# Single cell RNA sequence analysis of human bone marrow samples reveals new targets for isolation of skeletal stem cells using DNA-coated gold nanoparticles

**DOI:** 10.1101/2020.06.17.156836

**Authors:** Elloise Matthews, Stuart Lanham, Kate White, Maria-Eleni Kyriazi, Konstantina Alexaki, Afaf H. El-Sagheer, Tom Brown, Antonios G. Kanaras, Jonathan West, Ben D. MacArthur, Patrick S. Stumpf, Richard O.C. Oreffo

## Abstract

There is a wealth of data indicating human bone marrow derived stromal cells (HBMSCs) contain the skeletal stem cell (SSC) with the potential to differentiate along the stromal osteogenic, adipogenic and chondrogenic lineages. However, despite these advances, current methods to isolate skeletal stem cells (SSCs) from human tissues have proved challenging as no single specific marker has been identified limiting understanding of SSC fate, immunophenotype and the widespread clinical application of these cells. While a number of cell surface markers can enrich for SSCs, none of the proposed markers, alone, provide a platform to isolate single cells with the ability to form bone, cartilage, and adipose tissue in humans. The current study details the application of oligonucleotide-coated nanoparticles, spherical nucleic acids (SNAs), to rapidly isolate human cells using mRNAs signatures detected in SSCs in real time, to identify stem and progenitor skeletal populations using single cell RNA sequencing. Based on scRNA-seq of samples from 11 patients, this method was able to identify novel targets for SSC enrichment, which were assessed in a total of 80 patients. This methodology was able to isolate potential SSCs found at a frequency of <1 in 1,000,000 in human bone marrow, with a capacity for tri-lineage differentiation *in vitro*. The current approach provides new targets and a platform to advance SSC isolation, enrichment with significant therapeutic impact therein.

## INTRODUCTION

Cell fate in culture, tissues, and organisms can be followed using a variety of techniques, although cell phenotype information is typically limited to cell surface epitopes [1]. The detection of a specific mRNA responsible for the expression of a certain protein can be used to determine and characterise the phenotype of the cell. However, traditional methods to assess specific mRNAs such as in-situ hybridisation, northern blot, or quantitative-PCR necessitate cell fixation or lysis to isolate the RNA, resulting in loss of the cells for further experiments.

Bone has the capacity to regenerate, evidence of the presence of a stem cell in bone. While this regenerative capacity has long been recognized, the *in vivo* identity of the skeletal population has only recently been confirmed [2-4]. There is a wealth of data indicating human bone marrow derived stromal cells (HBMSCs) contain the skeletal stem cell (SSC) with the potential to differentiate along the osteogenic, adipogenic and chondrogenic lineages. The availability of a SSC pool has garnered significant interest across the regenerative medicine community, given the potential clinical application [5]. Despite this, current methods to isolate SSCs from human tissues remain challenging in the absence of a single specific marker for the SSC.

The inability, to date, to isolate a homogenous skeletal stem cell population has hampered understanding of SSC fate, immuno-phenotype and simple selection criteria; all limiting factors in the widespread clinical application of these cells. Although, a range of cell surface markers can enrich for SSCs [2, 6, 7], none of the proposed markers, alone, holds the potential to isolate a homogenous stem cell populations with the ability to form bone, cartilage, and adipose tissue in humans.

Gold nanoparticles (AuNPs) offer a unique material platform providing tuneable, readily synthesised nanoparticles with the potential for surface ligand functionalisation with, for example, oligonucleotides. These oligonucleotide functionalised AuNPs, termed spherical nucleic acids (SNAs), display enhanced stability, excellent cell uptake and a degree of resistance to enzyme degradation. Critically, the nanoparticles can be tailored to enhance selectivity and specificity towards the detection of intracellular targets such as RNA. We have recently developed a protocol using AuNPs to rapidly isolate human cells enriched in SSCs without affecting long-term cell viability [8], which has now been used in 80 different patient marrow samples.

In the current studies, we have harnessed SNAs to rapidly isolate live human cells using mRNAs signatures detected in skeletal cells in real time, to identify stem and progenitor skeletal populations using single cell RNA sequencing (Drop-Seq). Drop-Seq provides a single cell RNA sequencing platform that allows full transcriptome sequencing of upwards of tens of thousands of individual cells within a single experiment [9]. We have used Drop-Seq to generate mono-disperse nanolitre-sized droplets at >2 kHz for single cell capture. These droplets are stochastically loaded with single cells and, following cell lysis, poly-adenylated transcripts from individual cells are captured on functionalised micro-particles.

In this study, we have applied Drop-Seq to uncover the transcriptome signatures of enriched SSC populations and to subsequently identify transcripts that distinguish SSCs from other human bone marrow cell types. The current scRNA-seq data reveals candidate markers which highly complement the AuNP cell sorting approach to enrich SSC populations, validated through *in vitro* clonogenic assessment. We present *SPARCL1, Decorin (DCN), Transferrin (TF)* and *Caldesmon1 (CALD1)* as novel markers which serve as targets for the isolation of live SSC populations. The current approach provides new targets and a platform to advance SSC isolation and enrichment with significant therapeutic impact therein.

## METHODS

### Cell Culture

Bone marrow samples were obtained from haematologically healthy patients undergoing hip replacement surgery under local ethics committee approval local ethics committee approval (LREC194/99/1 and 18/NW/0231 and 210/01). In brief, bone marrow was washed in α-MEM medium and passed through a 70 μm cell strainer. Only samples intended for incubation with SNAs were treated with RosetteSep Human Granulocyte Depletion Cocktail (StemCell Technologies, Cambridge, UK) adding 100 µL to bone marrow resuspended in 5 ml α-MEM medium for 20 minutes. All samples were subjected to density centrifugation using Lymphoprep™ (Lonza). The buffy coat layer, containing bone marrow mononuclear cells was washed in basal medium (α-MEM containing 10% FBS and 100 U mL^-1^ penicillin and 100 µg mL^-1^ streptomycin; Lonza). Cells were subsequently either sorted using MACS or incubated with SNAs.

### Enrichment of CD45-CD146+ skeletal progenitor, CD144+ endothelial and CD144-/CD106+ pericyte cell populations

Whole bone marrow was diluted 1:1 with α-MEM (Gibco) digested with collagenase IV (20U/mL, Thermo Fisher, 17104019) for 30 min at 37 °C under continuous rotation. Subsequently, bone marrow mononuclear cells were isolated as described by Kanczler et al. 2019 (PMID: 30729460). Where indicated, magnetic cell sorting (MACS) was conducted to enrich or deplete cells. Up to a total of 1×10^8^ BM-MNCs were used for each separation. Cells expressing CD45 (Miltenyi Biotec, 130-045-801), CD146 (Miltenyi Biotec, 130-093-596), CD144 (Miltenyi Biotec, 130-097-857), CD106 (Miltenyi Biotec, 130-104-123 and 130-048-102) were isolated using LS columns (Miltenyi Biotec, 130-042-401) according to manufacturer’s instructions. Each of three patient samples were sorted into three distinct populations; CD45- /CD146+, CD144+ and CD144-/CD106+ cells.

### Stro-1 Enrichment of Colony Forming Units-Fibroblasts (CFU-Fs)

Marrow cells were initially incubated with blocking buffer (α-MEM, 10 % human serum, 5 % FCS and 1 % bovine serum albumin) and subsequently with primary Stro-1 antibody (undiluted hybridoma culture supernatant) [10]. Following washes in buffer (2 mM ethylenediaminetetraacetic acid (EDTA) and 1 % BSA in phosphate buffered saline), cells were incubated with magnetic bead-conjugated secondary antibody (Miltenyi Biotec). After further washes, Stro-1 positive cells were collected by magnetic-activated cell sorting (MACS) according to manufacturer’s instructions (Miltenyi Biotec).

### Synthesis of spherical nucleic acids (SNAs)

SNAs were manufactured as previously detailed [11] and sequences used detailed in Table 1. All sense strands were modified with FAM dye at the 5’ end and a thiol at the 3’ end. All flare strands were functionalised with a Cy5 dye at 5’ end. For dual-SNA selection, *SPARCL1* and *SOST* SNAs were modified with a JOE fluorophore instead of Cy5 at the 5’ end of flare strands. SNAs were added to the cell suspension in basal medium (at 20×10^6^ cells per ml) at 0.2 nM for 1 hour at 37 °C with rotation mixing. Cells were then washed in basal medium and resuspended in FACS solution prior to FACS sorting.

**Table 1.**
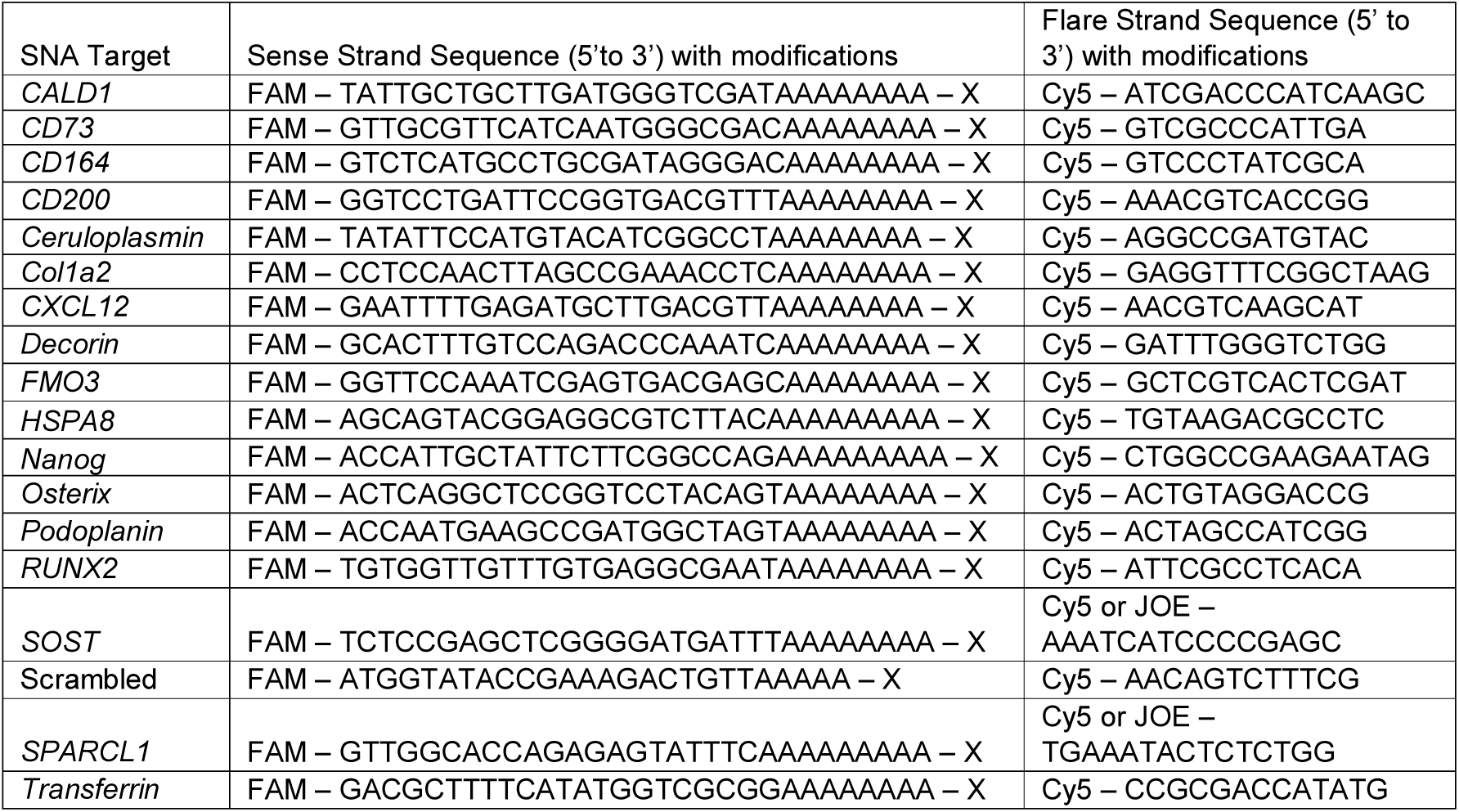
Sense and flare oligonucleotide sequences. X: thiol modifier 6 S-S (CPG resin from Glen Research)

### FACS

SNA treated cells were collected from human bone marrow using a FACS Aria cytometer (Becton Dickinson, Wokingham, UK). In total, 20 million cells were sorted from each patient sample. Cells were gated for monocytes, single cells, and Cy5 fluorescence, with the <u>top 20%</u> of fluorescent cells collected. Positive samples were determined to be within the top half of these Cy5 fluorescent cells and negative samples were considered to be within the bottom half of the collected cells. For dual-SNA studies, a four-way gating system sorted Cy5+/JOE+, Cy5-/JOE+, Cy5+/JOE-, Cy5-/JOE-cells, with only the top 10% of Cy5+/JOE+ collected for plating. Data were analysed on Flowing Software version 2.5 (http://www.flowingsoftware.com). Collected cells were subsequently used for CFU-F quantification, cellular differentiation assays, or Drop-Seq.

### Colony Forming Units-Fibroblast (CFU-F) Assay

CFU-F assessment was conducted to demonstrate colony growth [5, 12]. Depending on the number of cells collected by FACS, between 1,000 and 2,500 positive or negative FACS collected cells were placed into each well of 12 or 6-well tissue culture plates. Cells were grown for 14 days, with a medium change after 7 days. On day 14, wells were washed with PBS and then fixed with 95% ethanol for 10 minutes. The wells were then air dried and 1 mL 0.05% crystal violet solution added to each well for 1 minute. The wells were washed twice with distilled water and the number of stained colonies determined.

### Decorin and Transferrin Surface Antigen Detection

Surface antigen detection of Decorin and Transferrin was performed using MACS. Ten million granulocyte-depleted marrow cells were initially incubated with blocking buffer (α-MEM, 10 % human serum, 5 % FCS and 1 % bovine serum albumin) for 15 minutes at 4°C, then with primary antibody against Decorin (Abcam, ab181456) or Transferrin (Bio-techne, NBP2-02264) at 1/30 dilution for 30 minutes at 4°C. Following washes in buffer (2 mM ethylenediaminetetraacetic acid (EDTA) and 1 % BSA in phosphate buffered saline), cells were incubated with magnetic bead-conjugated secondary antibody (Miltenyi Biotec) for 15 minutes at 4°C. After further washes, MACS positive and negative cells were collected by magnetic-activated cell sorting (MACS) according to manufacturer’s instructions (Miltenyi Biotec). Cells were plated at 50,000 cells per well in a 12 well plate for MACS positive and negative cells and 5,000 cells per well for unsorted cells for CFU-F assessment.

### Osteogenic Differentiation Assay

Cells were cultured at 37 °C in 5% CO_2_ in T75 flasks until confluent. Cells were seeded at 10,000 cells per well on a 12 well plate. Cells were cultured in basal media for 24 hours. Cells were then cultured in osteoinductive media (basal medium with 50 µM ascorbic acid 2-phosphate and 10 nM vitamin D_3_) for 14 days at 37 °C in 5% CO_2_ with media change every 3-4 days. Cells were washed in PBS, fixed in 95% ETOH, stained with alkaline phosphatase.

### Adipogenic Differentiation Assay

Cells were cultured at 37 °C in 5% CO_2_ in T75 flasks until confluent. Cells were seeded at 10,000 cells per well on a 12 well plate. Cells were cultured in basal media for until 80% confluent, approximately 3-5 days. Cells were then cultured in adipogenic media (basal medium with 100 nM dexamethasone, 500 µM IBMX, 3 µg/ml ITS solution, and 1 µM rosiglitazone) for 14 days at 37°C in 5% CO_2_ with media change every 3-4 days. Cells were washed in PBS, fixed in 4% paraformaldehyde, washed in PBS again and subsequently stained with Oil Red O.

### Chondrogenic Differentiation Assay

Cells were cultured at 37 °C in 5% CO_2_ in T75 flasks until confluent. Cells were diluted to 500,000 cells per ml in chondrogenic media (α-MEM containing 100 U mL^-1^ penicillin and 100 µg mL^-1^ streptomycin, 100 µM ascorbic acid 2-phosphate, 10 ng/ml TGF-B_3_, 10 µg/ml ITS solution, 10 nM dexamethasone) in a universal container. Cells were centrifuged at 400 x g for 10 minutes to form a cell pellet, and all but 1 ml of media removed. Cells were then cultured with the tube cap loose for 14 days at 37 °C in 5% CO_2_ with media change every 2 days. Cells were washed in PBS, fixed in 95% ETOH, stained with alkaline phosphatase. Fixed pellets were stained with alcian blue and Sirius red.

### Microscopy

Cells were cultured at 37 °C in 5% CO_2_ in 24 well plates and imaged using a Zeiss Axiovert 200 inverted microscope with an Axiocam MR camera for fluorescent imaging and Axiovert HR camera for brightfield imaging operated by Zeiss Axiovision software version 4.7.

### Statistics

Wilcoxon-Mann-Whitney statistical analysis and ANOVA were performed where appropriate using the SPSS for Windows program version 23 (IBM Corp, Portsmouth, Hampshire, UK). Data is presented as mean ± 95% confidence limits and significance was determined with a p-level of 0.05 or lower.

### Drop-Seq of Human Bone Marrow Cells

Drop-Seq was performed as previously described [9] and outlined in the online Drop-Seq Laboratory Protocol version 3.1 (http://mccarrolllab.org/dropseq/) with any adjustments described below. Drop-Seq samples were processed as two separate experiments, with the first including the CD45-/CD146+, CD144+ and CD144- /CD106+ populations and the second including the SNA-Cy5+ and Stro1+ populations (which for the purpose of clarity, will be known as Drop-Seq1 and Drop-Seq2 respectively). In brief, droplet microfluidic devices, according to Macosko et al. [9] were fabricated in poly(dimethylsiloxane) (PDMS) and functionalised to an oleophilic state by incubating channels with 1% trichloro(1*H*,1*H*,2*H*,2*H*-perfluoro-octyl)silane (Sigma-Aldrich, 448931) in HFE-7500 (3M) for 5–10 min at RT. Cells were co-encapsulated with Drop-Seq beads (ChemGenes: lot 083117) in 1 nL droplets using 15000 mL/hr oil (QX200, Biorad), 4000 mL/hr cell suspension and 4000 mL/hr bead flow rates (droplet generation frequency 2 kHz). Following cell lysis, the droplet emulsion was broken using perflouroctanol (Sigma-Aldrich) to generate single-cell transcriptomes attached to microparticles (STAMPs). cDNA synthesis was conducted using Maxima H Minus Reverse Transcriptase (Thermo Fisher Scientific). This was followed by PCR amplification (Kapa HiFi Hotstart ReadyMix) using 4+15 PCR cycles for Drop-Seq1 and 4+14 cycles for Drop-Seq2 (95 °C 3 minutes - 4 cycles of: 98 °C 20 s; 65 °C 45 s; 72 °C 3min – 15/14 cycles of: 98 °C 20 s; 67 °C 20 s; 72 °C 3 min - 72 °C 5 min; 4 °C hold). Libraries were tagmented (Nextera XT DNA Library Preparation Kit, Illumina) and amplified before pooling samples for paired-end sequencing using a NextSeq 500/550 High Output Kit v2.5 (Illumina, 20024906) and NextSeq 500 system (Illumina).

### Sequence alignment

Raw sequencing reads were aligned following the Drop-Seq Core Computational Protocol (Drop-Seq tools v1.0, http://mccarrolllab.org/dropseq/). Sequencing data was aligned to human hg19 reference genome (GSM1629193) using STAR software [13]. Raw sequencing reads were demultiplexed, grouping reads by cell barcode to generate a digital gene expression (DGE) matrix for downstream analysis, using Drop-Seq tools (v1.0). A modified multi-mapper pipeline was executed to correct multiple alignment [14].

### Data clustering

Analysis of the Drop-Seq1 and Drop-Seq2 datasets was conducted using the software R (version3.5.0) according to the standard pipeline of functions included in the R Seurat package (version 3.0) (http://satijalab.org/seurat/). Drop-Seq1 and Drop-Seq2 datasets were processed and analysed separately.

#### Drop-Seq1

The Drop-Seq1 dataset, comprising 26,657 cells, was subjected to various quality control measures; filtering out of low-quality cells (cells expressing <200 genes or >10% mitochondrial genes) and potential cell doublets (cells expressing >4,500 genes). Cells were then normalised using the function; *sctransform*. Linear dimensionality reduction was performed using Principle Component Analysis (PCA) and non-linear dimension reduction using Uniform Manifold Approximation and Projection (UMAP). Clustering analysis was conducted using the Seurat function, *FindClusters*, at a resolution of 1.1 and default parameters. To assign identifies to the clusters, the function, *FindAllMarkers*, was performed to characterise significant cluster-specific gene expression for comparison against previously described biomarkers for bone marrow cell populations. This revealed 15 distinct cell types. Pericyte and endothelial cell clusters were defined by expression of *VWF* and *LEPR* respectively.

#### Drop-Seq2

For the Drop-Seq2 dataset, comprising 1,900 cells, quality control thresholds were more refined due to the reduced number of cells. Cells expressing <8.5% mitochondrial RNA, or <200/>2,600 unique reads were filtered from the dataset. Cells were then normalised using the function; *sctransform*. Linear dimensionality reduction was performed using Principle Component Analysis (PCA) and non-linear dimension reduction using Uniform Manifold Approximation and Projection (UMAP). Clustering analysis was conducted using the Seurat function, *FindClusters*, at a resolution of 1.1 and default parameters. This revealed 15 clusters which were broadly classified into 8 cell types. Skeletal stem cell cluster was defined by expression of *CXCL12, LEPR* and *SPARCL1*.

## RESULTS AND DISCUSSION

### Drop-Seq of whole bone marrow reveals SPARCL1 as a potential SSC marker

A sub-endothelial perivascular origin of the SSCs in bone marrow is widely acknowledged [3, 15-17]. To characterise the cellular heterogeneity within human adult bone marrow and to identify potential markers for SSC enrichment, scRNA-seq was performed, using the Drop-Seq methodology [9]. Three cell populations per patient were fractioned for sequencing; the previously described CD45-/CD146+ skeletal progenitor population [3], CD144+ endothelial cells [18] and CD144- /CD106+ pericytes [19]. In total, 9,795 cells were profiled, reduced to 7,385 following quality control processing. To further increase the cell pool for identification of skeletal progenitors, this data was combined with sc-RNAseq data for unsorted bone marrow populations from 3 patients [14](data available from ArrayExpress under accession number E-MTAB-8630), enabling comparison between depleted and unfractionated bone marrow. In total, the combined data, shown in figure 1A, comprised 17,527 cells with an average of 2,082 transcripts per cell.

**Figure 1:**
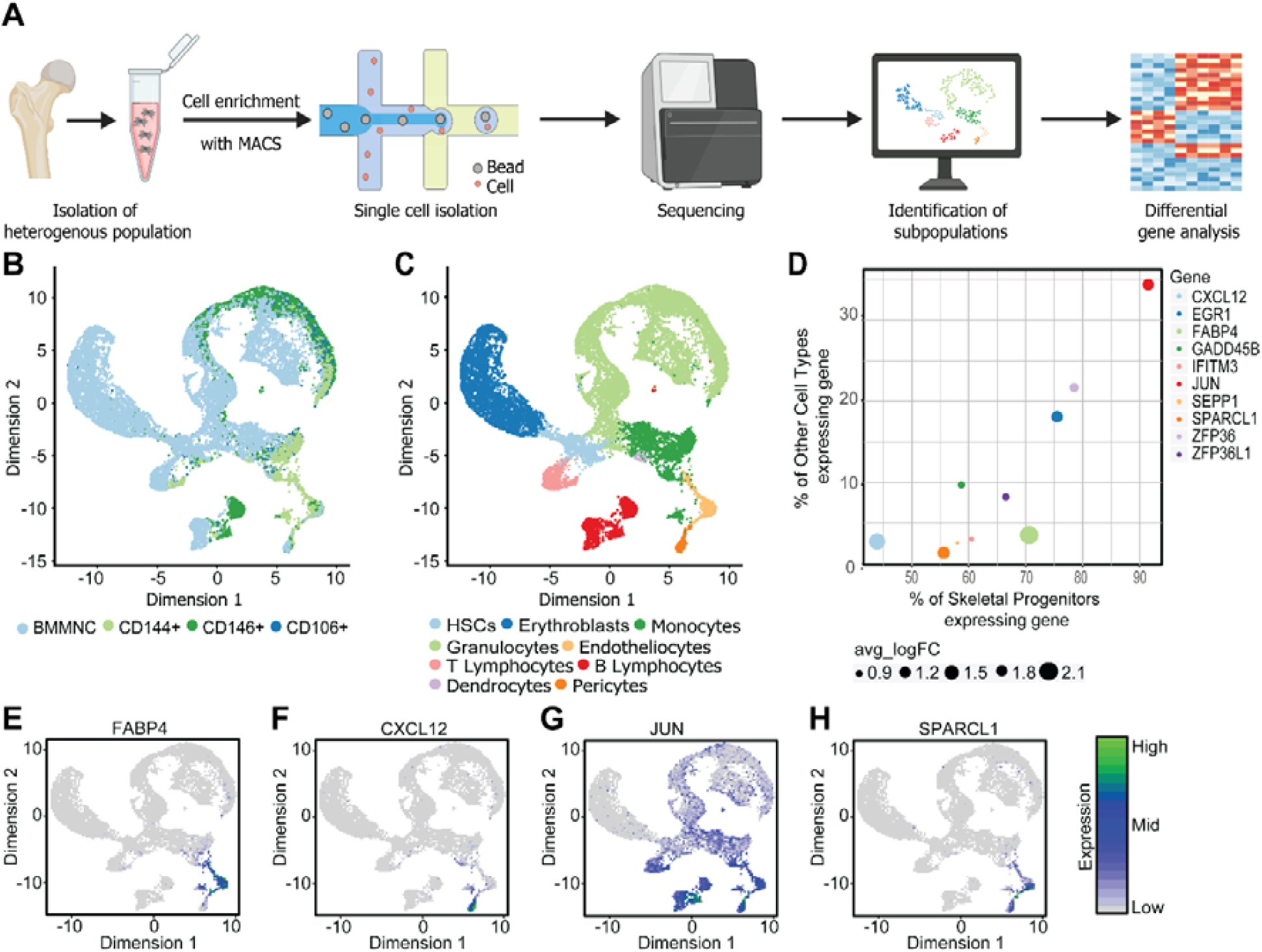
ScRNA-seq of 17,527 cells reveals *SPARCL1* as a candidate marker of skeletal progenitors. **A)** Experiment overview. Heterogeneous bone marrow populations were isolated from human adult bone marrow and MACS was applied to enrich for a CD45-/CD146+ skeletal progenitor population, CD144+ endothelial cells and CD144-/CD106+ pericytes. Single cells were isolated and sequenced following the Drop-Seq methodology. Unsupervised clustering was applied to reveal subpopulations which were characterised by differential gene expression to identify lineage biomarkers. **B)** Cell populations were mapped onto the UMAP representation to visualise spread of cells across clusters, each point representative of a single cell. **C)** Identification of lineage biomarkers characterised the clusters into 9 cell types. **D)** Top ten differentially expressed markers for skeletal progenitor clusters when compared to all other cell types. Embedding heatmap plots shown localised expression of **E)** *FABP4*, **F)** *CXCL12*, **G)** *JUN*, **H)** *SPARCL1*.

An unsupervised clustering strategy was employed to detect distinct cell subtypes with unique gene expression signatures [20]. Briefly, 9 cell clusters were identified, corresponding to hematopoietic and non-hematopoietic cell types (Figure 1B). Guided by the localised expression of established cell type markers, clusters were labelled as haematopoietic progenitors (*CD34*+, *SPINK2*+), granulocytes (*MPO+,LTF+*), erythroblasts (*HBD+, GYPA+*), B lymphocytes (*MZB1+, IGKC+*), monocytes (*CD14+, VCAN+*), T lymphocytes (*CD8A+, IL7R+*), dendrocytes (*FCGERA*+, *HLA-DQB2*+), endothelial cells (*VWF+, CLEC14A+*) and pericytes (*LEPR+, CXCL12+*) (Supplementary figure 1). We next sought to identify candidate SSC markers, through examination of localised gene expression in pericytes and endothelial cell clusters, representing 233 and 460 cells respectively. Analysis of differential gene expression between pericytes/endothelial cells, termed ‘Skeletal progenitors’, and other cell types indicated *SPARCL1*, as a potential SSC marker due to elevated expression (Figure 1C*). FABP4, CXCL12, JUN* and *SPARCL1* were among the 4 most differentially expressed genes in pericytes and endothelial cells (Figure 1D-G). However, from these mRNA, *SPARCL1* expression was strongly localised to the SSC region, with expression observed in 55.5% of skeletal progenitors and only 1.3% in other cell types. Given that *SPARCL1* has been previously shown to activate BMP/TGF-β signalling to promote the differentiation of C2C12 cells in mice [21], is highly homogenous to bone regulatory protein SPARC/OCN [22], new SNAs were designed against *SPARCL1* mRNA to investigate *SPARCL1* as a new target for SSC enrichment.

### SNAs targeting *SPARCL1, SOST, CD200* and *CD146* mRNA enrich for CFU-F

To test suitable candidate SNAs for SSC isolation, SNAs were designed targeting a range of mRNAs; these included SPARCL1, as identified by scRNA-seq, and other SNAs including: *HSPA8 (*encodes antigen detected by STRO-1 antibody)[23], *RUNX2* [24], and scrambled control. Other potential mRNA targets were *CD164, CD146* [3], *SP7* (Osterix) [24] *CXCL12* [25] and *Sclerostin (SOST)*[26]. Recently, Chan et al., 2018 and Debnath et al., 2018 have indicated likely candidate protein markers for SSCs [6, 27], and thus, SNAs were designed targeting *CD73, CD200*, and *PODOPLANIN (PDPN)*. Following incubation with the SNAs, positive and negative fractions were collected. Each SNA was tested against a minimum of 3 different patients, and up to 3 SNAs were tested for each patient. In total, bone marrow samples from 18 different patients were tested. To determine the clonogenic capacity of SNA-sorted populations, cells were plated at limiting dilution, whereby each fibroblastic colony formed, is derived from a single SSC or progenitor, termed CFU-F [28]. The normalised CFU-F counts as a percentage of the unsorted cells are presented in figure 2. Enrichment of CFU-F in comparison to scramble control and unsorted cells was observed in all positive fractions, while the negative fractions displayed minimal CFU-F.

**Figure 2:**
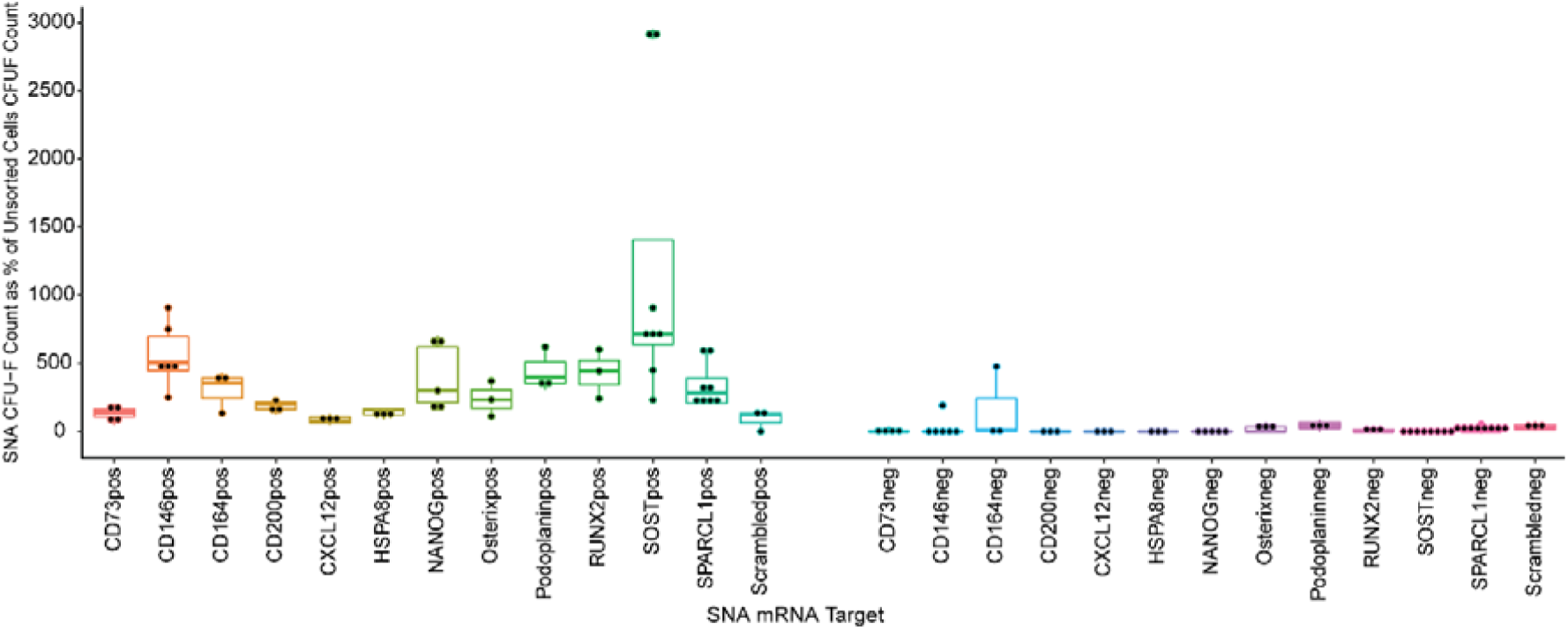
Cells enriched for *CD146, CD200, SOST*, and *SPARCL1* targets demonstrate enhanced CFU-F capacity in comparison to unselected cells. SNAs were designed to target a wide range of molecular markers, identified from scRNA-Seq libraries and related literature. The positive (red) and negative (green) populations, were plated at 5,000 cells per well of a 12-well plate and colonies were counted after two weeks of culture. For each SNA, each point represents the mean CFU-F count from a different patient, displayed as a percentage of unsorted CFU-F counts. Horizontal bars represent mean with SD error bars (blue).

The SNAs displaying the most consistent CFU-F enrichment, and specificity for the positive cell fraction in comparison to unsorted cells, were *CD200, SPARCL1, CD146, Nanog, SOST* and *osterix*. Interestingly, the Osterix SNA also demonstrated regular CFU-F enrichment in the negative cell fraction. In agreement with the scRNA-seq data, *SPARCL1 mRNA* facilitated sorting of cells with enhanced clonogenic functionality. The highest levels of CFU-F enrichment were observed in *SOST+* populations, presenting SOST as a candidate SSC biomarker. *SOST* is a negative regulator of *Wnt* signalling and consequently functions in a number of key pathways; including, stem cell proliferation [29], osteoblast activity [30] and mechanical response [31]. Elevated CFU-F enrichment was observed in *Nanog+* population, in accordance with the role of *Nanog* in stem cell maintenance [32, 33]. Similarly, *CD200+* and *CD146+* were defined as populations of interest for further scRNA-seq analysis. These results are consistent with previous findings which identify *CD200*+ [6, 34] and *CD146+* [3, 17, 35] as viable targets for SSC isolation. However, limitations to both populations have been identified; *CD200 mRNA* displayed poor SSC-specificity [36, 37], while SSCs have been identified in both *CD146+* and *CD146-/low* fractions [38, 39]. Therefore, these populations do not likely encompass a purified pool encompassing all SSCs, though these markers, will generate a valuable resource, together with *SPARCL1+* and *SOST+*, for further investigation and characterisation of SSCs.

### Sclerostin and SPARCL1 SNA display enhanced specificity for CFU-F formation compared to traditional Stro1+ MACS populations

Following the enrichment in CFU-F counts, studies were undertaken to evaluate if the SOST+ and *SPARCL1+* SNA-selected cells displayed enhanced CFU-F enrichment compared to the current SSC enrichment methodology using MACS to select Stro-1+ cells (experiments outlined schematically in figure 3A). BMMNCs were divided into two parts and processed separately. The first fraction was incubated with SNAs targeting *SOST* or *SPARCL1 mRNA* and cells were collected using FACS. Meanwhile, the remainder of the sample was incubated with Stro-1 antibody and Stro-1+ cells were collected using MACS methodology [40]. The Stro-1+ cells were sequentially sorted using the *SOST* or *SPARCL1* SNA to collect Stro1+*SPARCL1*+ or Stro1+*SOST+* populations. Positive and negative fractions were plated to determine CFU-F enrichment (Figure 3A). Evaluation of data from patients B and C (Figure 3B) demonstrated *SPARCL1+* SNA-selected cells produced greater numbers of CFU-F than unsorted and *SPARCL1-*fractions (black bars), although this did not reach significance. Additionally, no significant difference was observed between *SPARCL1+* SNA-selected cells and Stro-1+ MACS-only collected cells for patients A and C, indicating *SPARCL1+* cells contain the same number of CFU-F as Stro-1+ cells. It was previously shown that all CFU-F reside within Stro-1+ fraction [41], therefore the current study suggests *SPARCL1* is a comparable target for SSC enrichment, with the additional benefits of the SNA cell isolation protocol being less time-consuming and therefore advantageous. Furthermore, when Stro-1+ were subsequently sorted using *SPARCL1* SNAs, significantly enhanced CFU-F enrichment was observed in Stro-1+*SPARCL1+* cells compared to Stro1+*SPARCL1-* (p<0.001 for all patients) and *Stro-1+* (only) (p≤0.03 for all patients) populations. These findings demonstrate the *SPARCL1* SNA holds enhanced specificity for CFU-Fs selection in comparison to Stro-1 antibody.

**Figure 3:**
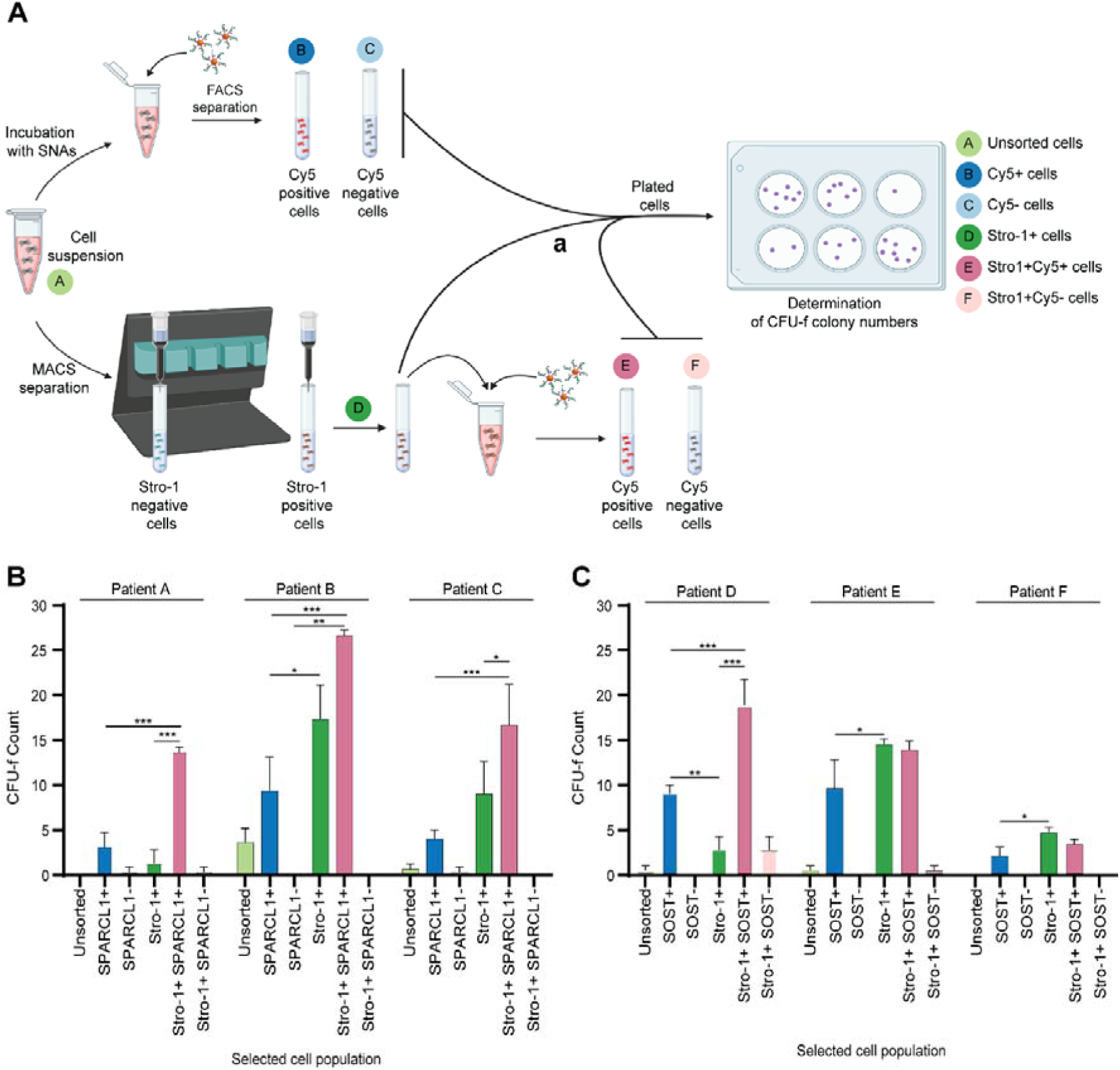
Comparison of SPARCL1 and SOST SNA to Stro-1 antibody for CFU-F isolation. A. Schematic of experimental protocol for different cell populations collected. Cells were either unselected (black bars) or Stro-1+ selected cells (green bars) from the same patient sample. B). Using SNA targeting *SPARCL1*. C). Using SNA targeting SOST. Only selected statistics are shown for clarity. * p<0.05, *** p<0.001. Bars show mean with 95% confidence limits. n=3 for each timepoint for each patient.

For all patients, *SOST+* SNA-selected cells produced greater numbers of CFU-F than unsorted and *Sclerostin-*fractions (black bars, all p<0.05) (Figure 3C). Patient D showed significantly enhanced CFU-F formation in the *Sclerostin+* SNA-selected cells than the Stro-1+ MACS-only collected cells (p<0.01). In contrast, for patients E and F the CFU-F levels observed were lower in the *Sclerostin+* SNA-selected cells (p<0.05). When Stro-1+ were subsequently sorted using *Sclerostin* SNAs, significantly enhanced CFU-F enrichment was observed in Stro-1+*Sclerostin+* cells compared to Stro1+*SPARCL1-* (p=0.001 for all patients) and Stro-1+ (only) (p<0.001 for patient D) populations. For patients E and F, the Stro-1+ CFU-F were not significantly different, indicating the *Sclerostin* SNA also holds enhanced or equal specificity for CFU-Fs compared to Stro-1 antibody.

### Drop-Seq of SNA-selected populations reveal novel targets for SSC isolation

The scarcity of SSC within human bone marrow, <1 in 100,000 of bone mononuclear cells [42], presents a significant obstacle for transcriptomic profiling of SSCs, as limitations of scRNA-seq protocols in detecting cell types with a frequency less than 1% have been recently reported as these cells typically appear as outliers [43]. Therefore, pre-sorting cells prior to downstream scRNA-seq can increase read depth by reducing competition for sequencing capacity between cells and genes [44]. Sequencing of populations enriched for desirable cell types, has been applied previously for the characterisation of rare cell subtypes and identification of unique biomarkers [45, 46]. In order to increase the SSC pool for scRNA-seq, Drop-Seq of MACS-collected Stro-1+ cells, and cells enriched using SNAs targeting *CD146, CD200, SOST* and *SPARCL1 mRNAs* was performed (Figure 4A). Though *Nanog* offers a promising SNA target for SSC enrichment, we were unable to collect enough *Nanog+* cells to obtain a sufficient library for scRNA-seq. In total the transcriptomes of 1,900 cells were sequenced, producing a dataset of 1,512 cells with an average of 1,956 transcripts per cell following quality control filtering. Unsupervised clustering produced 15 clusters which, following analysis of differential gene expression, were classified as 8 distinct cell types; Skeletal stem cells/progenitors, monocytes, erythroblasts, granulocytes, B lymphocytes, dendritic cells, T lymphocytes and Haematopoietic stem cells (Figure 5B). For example, clusters 0, 3, 6 and 11 showed high expression of the markers; *Glycoprotein-A* (GYPA) and Haemoglobin subunit alpha 1 (HBA1) (Supplementary figure 2), therefore identifying these cell clusters as erythroblasts.

**Figure 4:**
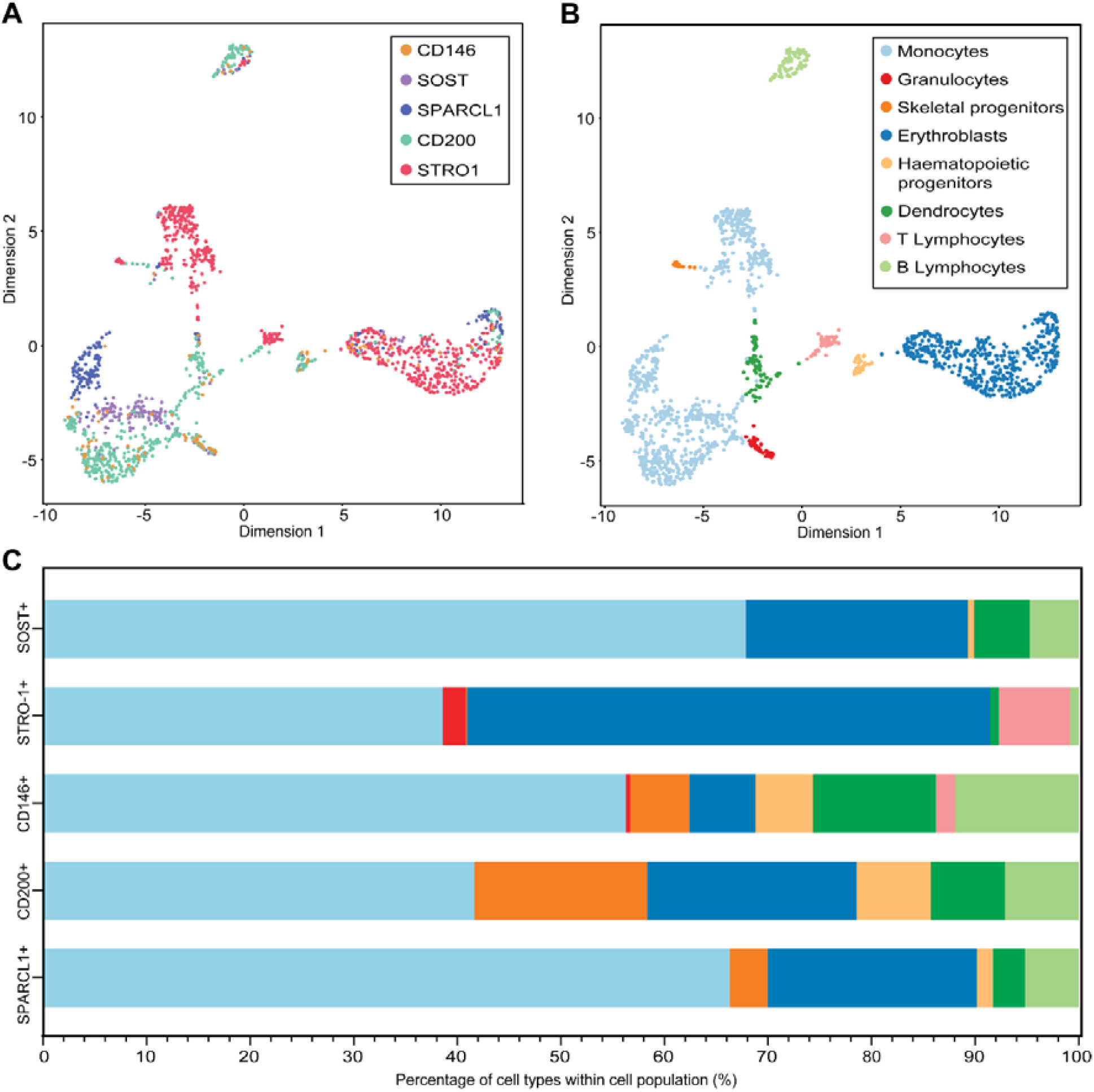
Drop-Seq analysis of 1,512 selected HBMSCs. UMAP was conducted for visualisation to reduce the data to two dimensions. Each point represents a single cell. A) Cells were sorted using SNAs targeting CD146, CD200, *SOST* and *SPARCL1* and sequenced, together with a MACS Stro-1+ population, following the Drop-Seq methodology. B) Unsupervised clustering at a resolution of 1.3 identified 15 subpopulations based on gene expression profiles. Clusters were classified into 8 distinct cell subpopulations, corresponding to haematopoietic and non-haematopoietic cell types. C) The proportion of cells from each sequenced population represented across the cell-type clusters reveals enrichment of monocytes within SNA-sorted populations (*CD200*+, *CD146*+ *SOST*+ and *SPARCL1*+). The Stro1+ population is predominantly erythroblasts and monocytes.

**Figure 5:**
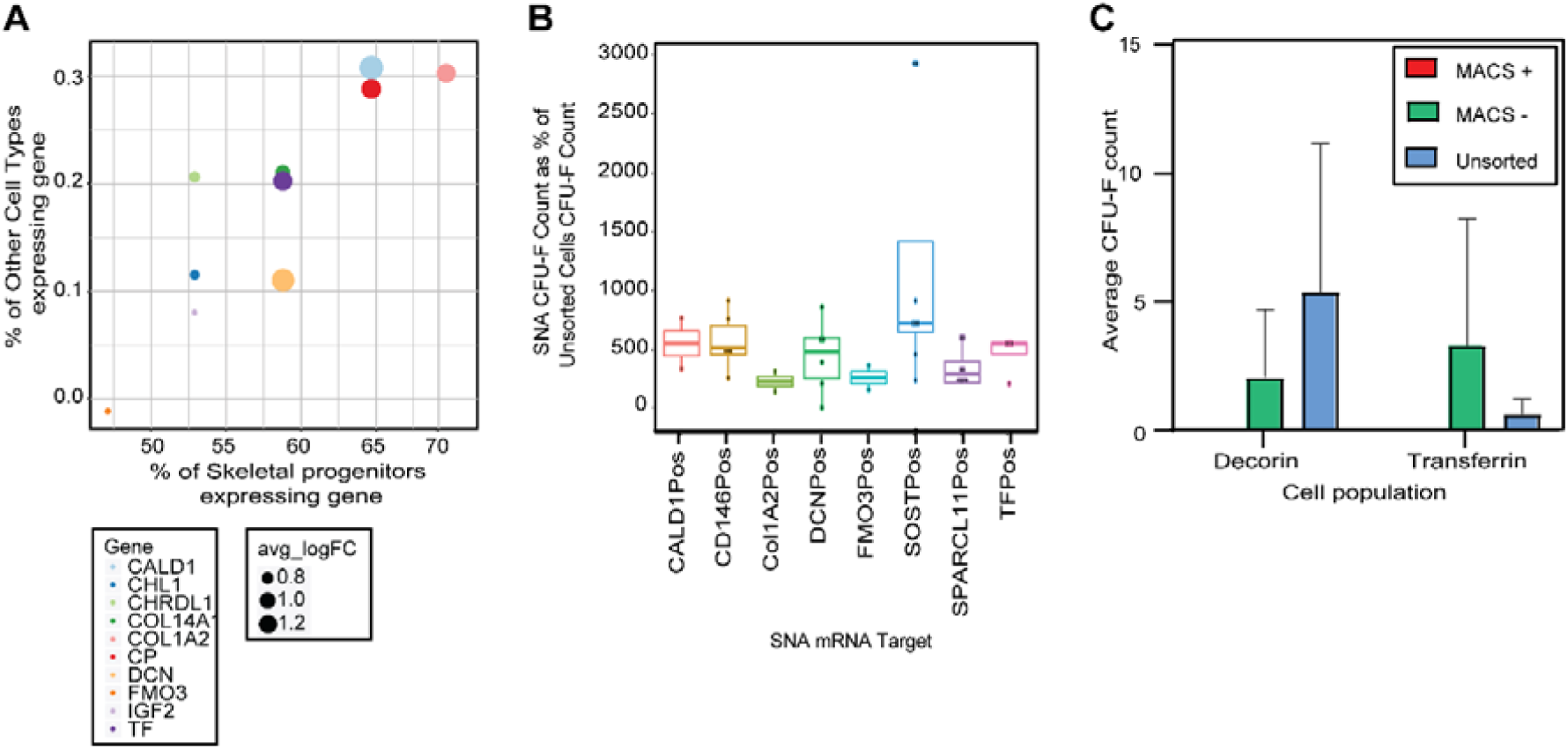
**A) Proportion of ‘Skeletal Progenitors’ and ‘Other cell types’ expressing the top 10 differentially expressed genes.** The ten most differentially expressed genes were plotted, with data-point size indicative of the average logarithm fold change (avg_logFC) in gene expression. **B) SNAs were designed to target *DCN, TF, CALD1, COL1A2* and *FMO3*; markers identified following Drop-Seq of Stro-1+, *SOST+, SPARCL1*+, *CD200*+ and *CD146*+ populations.** The positive and negative populations, together with unsorted cells, were plated at 5,000 cells per well of a 12-well plate and colonies were counted after two weeks of culture. For each SNA, each point represents the mean CFU-F count from a different patient, displayed as a percentage of unsorted CFU-F counts. In total, bone marrow samples from 17 different patients were tested. Horizontal bars represent mean with SD error bars. **C) Average CFU-F count for positive and negative fractions for DCN and TF MACS collected selected populations, together with CFU-F counts in unsorted cells of the same samples.** MACS was performed to sort HBMSCs based on expression of DCN and TF. Positive and negative fractions, together with unsorted HBMSCs, were plated. After two weeks of culture, colonies were stained and CFU-F counted.

Skeletal progenitor identities were assigned based on expression of stromal markers; *CXCL12* [25] and *LEPR* [47], and the expression of *SPARCL1*. The skeletal progenitor cluster consisted of 12.5% *SPARCL1*+ cells and 87.5% Stro-1+ cells, supporting the presence of SSCs within these enriched populations. No cells belonging to the *CD146*+, *SOST+* and *CD200*+ sequenced populations were represented within the cluster of skeletal progenitors. Somewhat surprisingly, *SOST* transcripts were not detected in any cells. Overall, the sequenced populations maintained a degree of heterogeneity and included cells across the classified cell-types; *SPARCL1*+ cells were represented in all 8 cell-type subpopulations (Figure 4C). Despite this observed heterogeneity, the SNA methodology was found to enrich for monocytes; with *CD146+, CD200+, SOST+* and *SPARCL1*+ populations consisting of 56%, 42%, 68% and 66% monocytes, respectively (Figure 4C), therefore demonstrating the efficiency of the SNA protocol in filtering and selection of cell types. Similarly, the MACS Stro-1+ population contained a high proportion of monocytes (39%), though Stro1+ cells were largely erythroblasts (51%), consistent with the findings of Simmons and Torok-Strob who demonstrated that CFU-F predominantly resided within the Stro-1+/GYPA-fraction [41].

### Novel SSC targets; *DCN, TF* and *CALD1* show consistent enrichment of CFU-F and enhanced enrichment of CFU-F in comparison to *SPARCL1+* cells

To further characterise the molecular signature of SSCs, differential gene analysis was performed to identify markers with localised expressed in skeletal progenitor cell types. The top ten differentially expressed genes in the SSC clusters are shown in figure 5A. These genes represent potential candidate SSC markers given the extremely low representation observed within other cell types; >0.75%. The most highly expressed differential marker was *Decorin* (*DCN*), expressed by 63% of skeletal progenitors with an avg_LogFC in gene expression of ∼1.5. In addition to *DCN*, 4 other markers; *Collagen Type I Alpha 2 Chain (COL1A2), Flavin-containing monoxygenase-3* (*FMO3), Transferrin (TF)* and *Caldesmon (CALD1)* were identified as potential candidates for SSC enrichment based on exclusivity and level of expression in skeletal progenitors compared to other cell types (figure 5A).

To assess the functionality of the novel markers, for isolation of SSCs, SNAs targeting *DCN, TF, COL1A2, CALD1*, and *FMO3* were designed. HBMSCs were incubated with the SNAs for 1 hour before positive and negative Cy5 cell fractions were sorted. SNAs were tested on a minimum of two different patient samples per target. For each sample, positive, negative and unsorted cells were plated at 5,000 cells per well of a 12 well plate to assess SSC enrichment within each fraction. Following crystal violet staining, visible clusters were counted, indicative of the number of SSC colonies. The number of CFU-F for each selected population was expressed as a percentage of unsorted cells (Figure 5B). Furthermore, the CFU-F value for *SPARCL1*+ and *CD146+* cells (previously showed the highest CFU-F enrichment), was included to allow comparison with the new SNA populations. For each SNA-target, enrichment for CFU-F was observed in all positive fractions. *DCN+, TF+* and *CALD1+* cells showed enhanced enrichment in comparison to *SPARCL1+* cells and comparable enrichment to *CD146+* cells, and therefore defined as candidate SSC biomarkers. A role of *DCN* in skeletal development has been identified previously; high expression has been reported in a variety of cells, including osteoblasts, perivascular chondrocytes and throughout the bone periosteum [48]. Additionally, DCN can function in signalling cascades; regulating cell differentiation, angiogenesis and bone mineralisation [49-54]. Similarly, the essential role of *TF* in iron-delivery and consequently, cell growth and survival, correlates with highly proliferative cell types and anti-apoptotic activity [55-58]. Finally, *CALD1* functions in calcium-mediated signalling pathways in bone [59], and its expression in BMMNCs has been reported previously [60]. In summary, there is emerging evidence to substantiate *DCN, TF* and *CALD1* as useful targets to explore for SSC isolation.

MACS provides a less time-consuming approach to cell sorting of large cell numbers than the SNA-based FACS strategy applied in this study, although MACS relies on the detection of surface epitopes [61]. It was therefore of interest to investigate whether the same level of SSC enrichment, observed following SNA-based cell sorting, could be achieved using MACS, with *DCN* and *TF* as antigen targets. Negative and positive fractions were collected for three samples and cells were plated at 50,000 cells per 3 wells of a 12-well plate for CFU-F assessment. In marked contrast to the CFU-F enrichment observed in *DCN+* and *TF+* populations following AuNP mRNA detection, cells sorted using DCN and TF antibodies did not display CFU-F capacity in positive fractions following MACS (figure 5C). This result indicates the molecular targets identified by scRNA-seq for SNA cell sorting are not suitable for surface epitope-based strategies, confirming mRNA levels do not directly correlate into protein synthesis [62-64]. This limitation was previously described by Fitter et al who identified no correlation between *HSPA8* (mRNA) and Stro-1 (surface protein) [23].

### Dual-SNA incubation collects cell populations with the capacity for CFU-F enrichment and tri-lineage differentiation

Following identification of SNA targets that sort populations with an enhanced CFU-F capacity, 2^nd^ generation dual SNAs targeting *SOST* or *SPARCL1 were* designed using JOE fluorescent dye in the place of Cy5 on the 5’ end of flare strands. The choice of dye made no difference to the CFU-F count isolated (data not shown). Cells were incubated with JOE labelled SNAs targeting either *SPARCL1* or SOST, together with Cy5 labelled SNAs targeting either *CD146, CALD1, COL1A2, DCN, FMO3, Nanog* or *TF*. The top 10% Cy5+/JOE+ fluorescent cells were collected and plated for CFU-F assessment. *SPARCL1+TF+, SPARCL1+Nanog+ and SPARCL1+CD146+* cells demonstrated the most enhanced CFU-F enrichment compared to unsorted cells (figure 6a).

**Figure 6.**
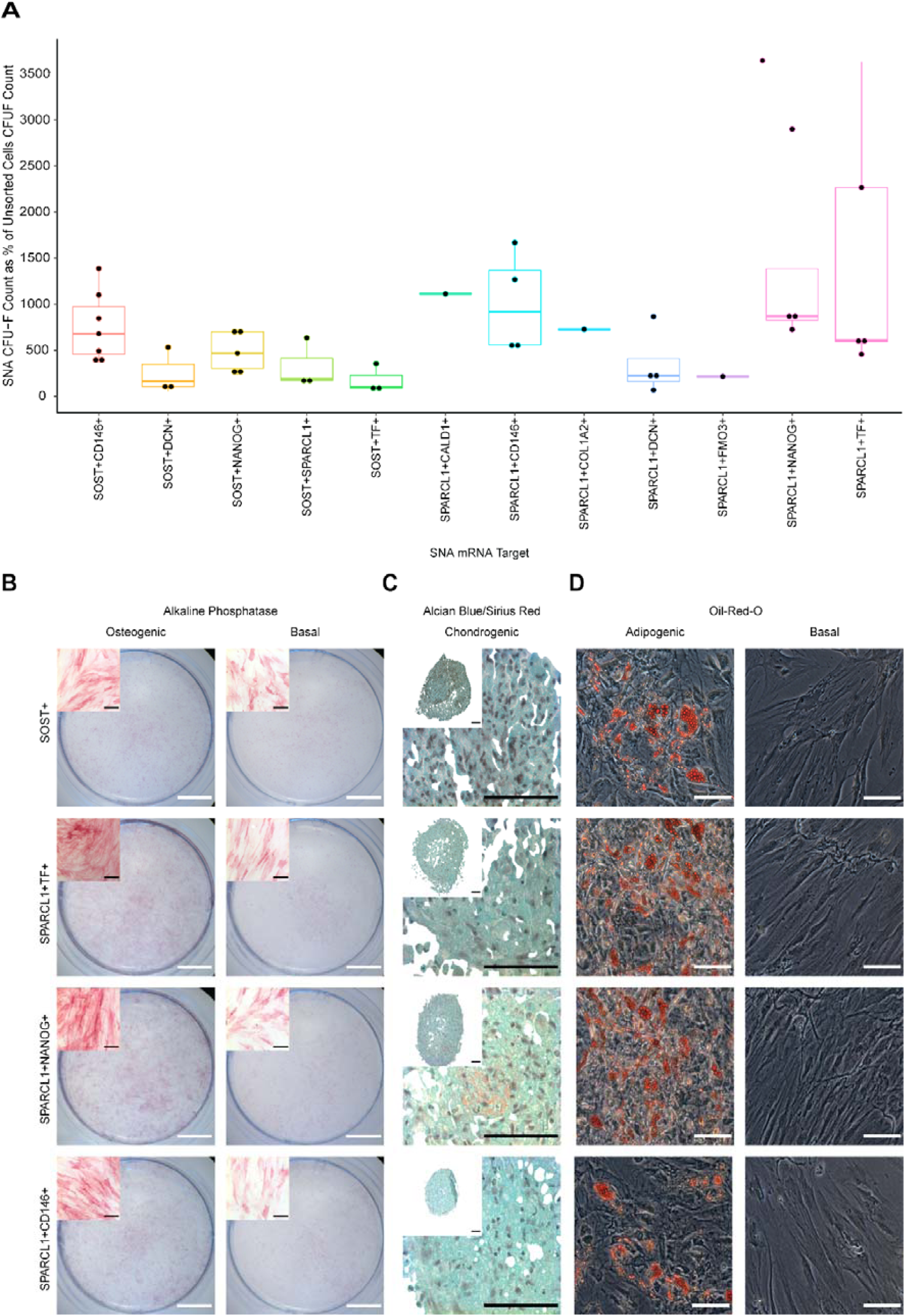
Dual-SNA selected cell populations display enhanced levels of CFU-F and capacity for tri-lineage differentiation. **A) CFU-F following dual-SNA selection.** Cells were incubated with SOST or SPARCL1 JOE-tagged AuNPs and Cy5-tagged AuNPs targeting *CD146, CALD1, DCN, COL1A2, FMO3, Nanog* or *TF*. The positive and negative fluorescent cells were collected and plated for CFU-F. For each SNA combination, each point represents the mean CFU-F count from a different patient, displayed as a percentage of unsorted CFU-F counts. In total, bone marrow samples from 18 patients were tested. Horizontal bars represent mean with SD error bars. *SPARCL1+CD146+, SPARCL1+Nanog+, Sparcl1+TF+ and SOST+* populations were collected and expanded in vitro. **B) Osteoinduction.** Cells were cultured in basal medium with 50 µM ascorbic acid 2-phosphate and 10 nM vitamin D_3_ for 14 days. Mineralisation is shown using alkaline phosphatase staining. Scale bars =100 µm, whole well = 500 µm. n=3 **C) Chondrogenic induction.** Cells were cultured in basal medium supplemented with 100 µM ascorbic acid 2-phosphate, 10 ng/ml TGF-B_3_, 10 µg/ml ITS solution, 10 nM dexamethasone. Alcian blue/Sirius Red staining revealed proteoglycan synthesis (Blue denotes proteoglycan deposition, purple indicates collagen deposition). Scale bars = 100 µm. n=3. **D) Adipogenic induction.** Cells were cultured in basal medium with 100 nM dexamethasone, 500 µM IBMX, 3 µg/ml ITS solution, and 1 µM rosiglitazone for 14 days. Oil-Red-O staining indicates lipid droplet formation. Scale bars = 100 µm. n=3.

Across all CFU-F studies, the SNAs that collected cells displaying the highest levels of CFU-F enrichment were SOST+ (Figure 2) and *SPARCL1+TF+, SPARCL1+Nanog+* and *SPARCL1+CD146+* dual SNAs (Figure 6a). To assess the stem cell potential of these subpopulations, we expanded SNA-enriched SOST+, *SPARCL1+TF+, SPARCL1+Nanog+* and *SPARCL1+CD146+* populations *in vitro* and cultured cells under conditions favourable for induction of SSC tri-lineage differentiation, an established criteria defining SSCs [42]. For osteoinduction, cells were cultured in basal medium with 50 µM ascorbic acid 2-phosphate and 10 nM vitamin D_3_ for 14 days. Histological analysis revealed enhanced expression of alkaline phosphatase in SNA-sorted populations following culture in osteogenic media in comparison to basal cultured populations. Positive staining for alkaline phosphatase was observed in basal-cultured populations though at a reduced level, inferring an elevated osteogenic capacity in the SNA-sorted populations (Figure 6b). For chondrogenic differentiation assessment, cells were cultured as pellets in basal medium with 100 µM ascorbic acid 2-phosphate, 10 ng/ml TGF-B_3_, 10 µg/ml ITS solution, 10 nM dexamethasone. Alcian blue staining of the chondrogenic pellets revealed proteoglycan synthesis in all SNA-sorted populations. However, Sirius Red stain retention was absent in all pellets, except minimal staining observed in *SPARCL1+Nanog+* cells, indicating no collagenous matrix formation (Figure 6c). To evaluate the adipogenic potential of the SNA-sorted populations, cells were cultured in 100 nM dexamethasone, 500 µM IBMX, 3 µg/ml ITS solution, and 1 µM rosiglitazone. Oil-Red-O staining of lipids was performed after 14 days of culture, evidencing induction of adipogenesis and lipid droplet formation, with enhanced levels of lipid droplets observed in *SPARCL1+Nanog*+ and *SPARCL1+TF*+ populations (Figure 6d). In summary, we evidence the enhanced proliferation and multi-lineage potential of SNA-sorted populations following culture in supplemented media, therefore fulfilling the criteria of SSCs and confirming the prospective applications of SNAs within the field of skeletal regeneration.

## CONCLUSION

In summary, the current studies demonstrate single cell RNA sequencing provides a powerful tool for the parallel single-cell sequencing of human adult bone marrow utilizing DNA-coated gold nanoparticles. This approach revealed significant heterogeneity within unsorted and enriched bone marrow populations. Initial profiling of pericyte, endothelial, skeletal progenitor and unsorted cells revealed *SPARCL1+* as a potential SSC biomarker. This profiling was only possible due to the ability of the protocol used to isolate rare cell populations. Thus, the Drop-Seq data using *CD146, CD200, SOST* and *SPARCL1* SNAs, together with a Stro-1+ population, produced transcripts from 1,512 cells form an initial 200 million bone marrow cells. When targeted using AuNP mRNA detection, *SPARCL1+* cells displayed a 3-fold enhanced capacity for CFU-F in comparison to unsorted cells and improved specificity for CFU-F enrichment in comparison to the widely accepted Stro-1+ pool of SSCs [41]. Similarly, *CD146* and *CD200* were identified as mRNA targets for AuNP cell sorting to enrich CFU-F in SSC populations. Delivery of *SPARCL1*+, *CD146*+, *CD200*+, and Stro-1+ populations into the Drop-Seq methodology revealed a second generation of markers, which demonstrated improved capacity for CFU-F enrichment. *DCN, TF* and *CALD1* were identified as novel markers that increased CFU-F capacity when used both alone or in conjunction to *SPARCL1-*targeted SNAs.

Although *Sclerostin* showed initial potential for CFU-F enrichment, the lack of transcript detection in the Drop-Seq cells, together with an inability to demonstrate for CFU-F enrichment in conjunction with CD146, DCN or TF SNAs potentially suggests the SNA sequences may not be specific for Sclerostin, despite showing specificity on BLAST searches. In addition, unlike *SPARCL1* which showed a consistent pattern compared to Stro-1 isolated cells from different patients, Sclerostin CFU-F enhancement was patient specific.

The current studies demonstrate, for the first time, that Drop-Seq can reveal novel markers for SSC isolation, which sort cells with clonogenic and multi-lineage differentiation functionality *in vitro*. Furthermore, we demonstrate scRNA-seq approaches identify markers that highly complementary for cell sorting strategies harnessing mRNA detection, like the AuNP protocol applied in this study, in contrast to epitope expression. Overall, the reported findings describe novel markers that enable the isolation of an enriched pool of SSCs from human adult bone marrow, a valuable resource for future development of SSC-based therapies in the treatment of skeletal damage and disease.

## Acknowledgements

ROCO and AGK acknowledge financial support from the Biotechnology and Biological Sciences Research Council (BB/P017711/1). ROCO acknowledges support from the UK Regenerative Medicine Platform “Acellular / Smart Materials – 3D Architecture” (MR/R015651/1), the Rosetrees Trust, Wessex Medical Research as well as Institute for Life Sciences and University of Southampton. PSS, BDM and JJW acknowledge funding from the Medical Research Council (MC_PC_15078).

---

**Supplementary figure 1.**
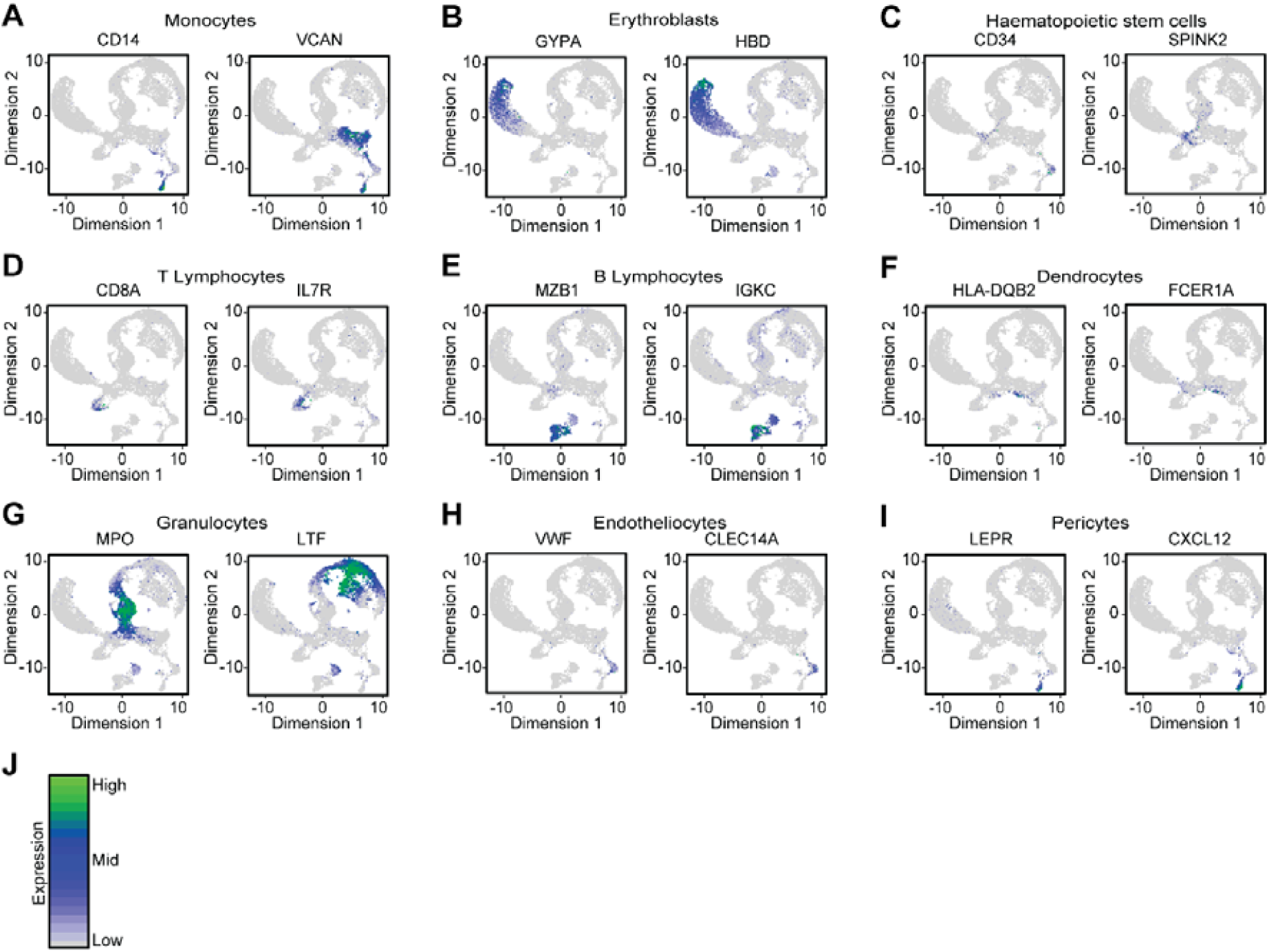
Use of lineage biomarkers to characterise Drop-Seq data clusters from CD45-/CD146+ skeletal progenitor population, CD144+ endothelial cells, CD144-/CD106+ pericytes and unselected bone marrow cells into 9 cell subpopulations.

**Supplementary figure 2.**
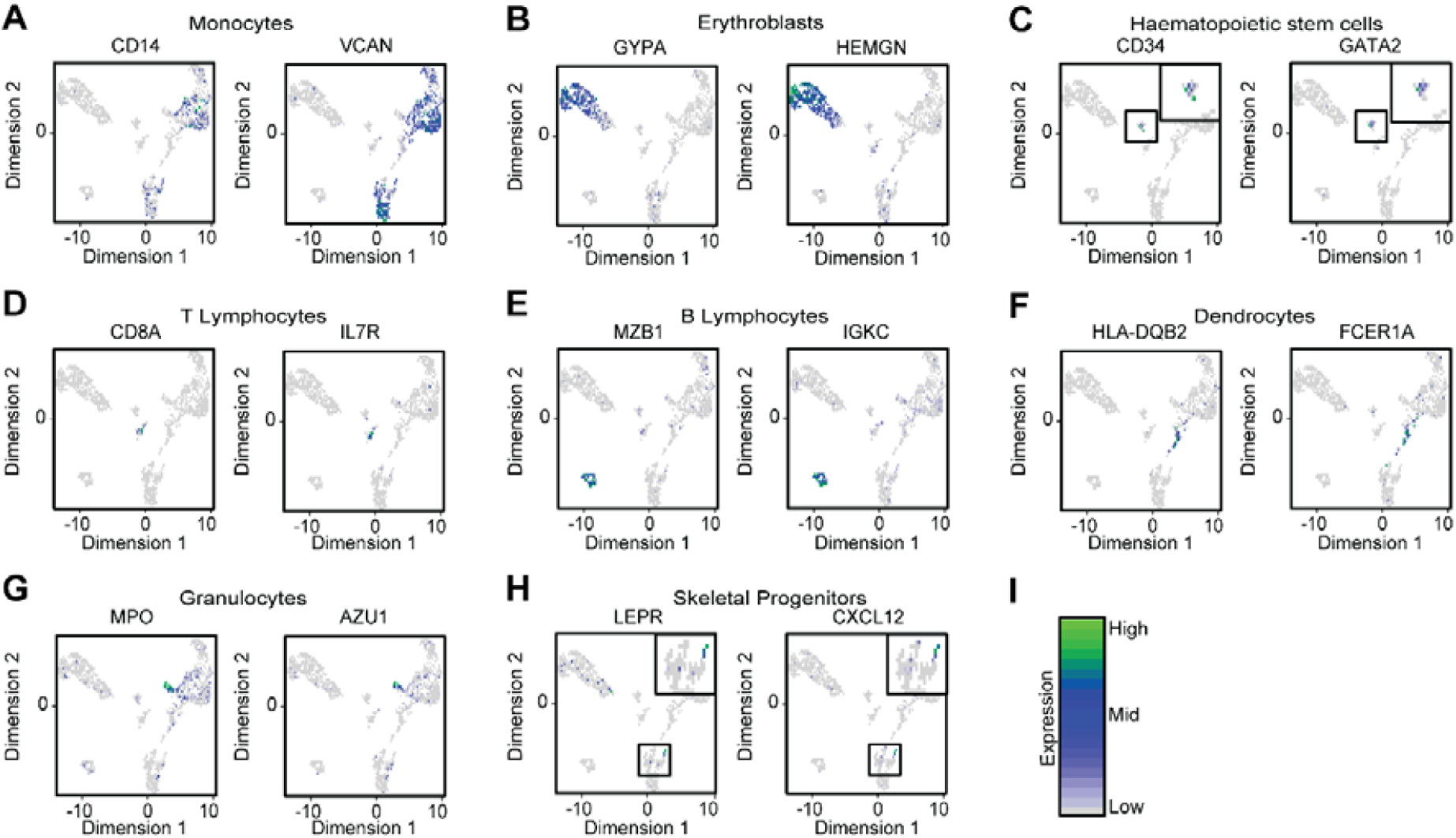
Use of lineage biomarkers to characterise Drop-Seq data clusters from Stro-1+ cells, and cells enriched using SNAs targeting CD200, SPARCL1 and CD146 mRNAs into 8 cell types.

## Notes

### Competing Interest Statement

The authors have declared no competing interest.

